# A quick and versatile protocol for the 3D visualization of transgene expression across the whole body of larval *Drosophila*

**DOI:** 10.1101/2021.01.28.428398

**Authors:** Oliver Kobler, Aliće Weiglein, Kathrin Hartung, Yi-chun Chen, Bertram Gerber, Ulrich Thomas

## Abstract

Larval *Drosophila* are used as a genetically accessible study case in many areas of biological research. Here we report a fast, robust and user-friendly procedure for the whole-body multifluorescence imaging of *Drosophila* larvae; the protocol has been optimized specifically for larvae by systematically tackling the pitfalls associated with clearing this small but cuticularized organism. Tests on various fluorescent proteins reveal that the recently introduced monomeric infrared fluorescent protein (mIFP) is particularly suitable for our approach. This approach comprises an effective, low-cost clearing protocol with minimal handling time and reduced toxicity in the reagents employed. It combines a success rate high enough to allow for small-scale screening approaches and a resolution sufficient for cellular-resolution analyses with light sheet and confocal microscopy. Given that publications and database documentations typically specify expression patterns of transgenic driver lines only within a given organ system of interest, the present procedure should be versatile enough to extend such documentation systematically to the whole body. As examples, the expression patterns of transgenic driver lines covering the majority of neurons, or subsets of chemosensory, central brain or motor neurons, are documented in the context of whole larval body volumes (using nsyb-Gal4, IR76b-Gal4, APL-Gal4 and mushroom body Kenyon cells, or OK371-Gal4, respectively). Notably, the presented protocol allows for triple-color fluorescence imaging with near-infrared, red and yellow fluorescent proteins.

## Introduction

Our understanding of the principles of animal form and function has considerably advanced in recent years. The combination of novel approaches to genetic manipulation with new techniques in sample preparation and microscopy, along with smart solutions in the processing of large image data sets, has indeed revealed much detail about the organization of various cell types, tissues and organs. What is missing, however, are techniques for the convenient, routine contextualization of such data within the whole body. Here, we report such a technique for larval *Drosophila.*

*Drosophila* larvae are used to study processes as diverse as chromatin organization and remodeling (Schwartz & Cavalli, 2017), tissue morphogenesis and regeneration (Heller & Fuchs, 2015; Vizcaya-Molina et al., 2020), tumorigenesis (Tamori & Deng, 2017), wound healing (Tsai et al., 2018) and innate immune responses (Buchon et al., 2014). Moreover, larval *Drosophila* serve as a model system in both the neural and the behavioral sciences. Studies on the third-instar larval neuromuscular junction (NMJ), for instance, have uncovered evolutionarily conserved mechanisms of synaptogenesis, synaptic function, and synaptic plasticity (Frank et al., 2020; Ghelani & Sigrist, 2018; Harris & Littleton, 2015). Current behavioral research is aimed at the characterization of the neural circuits underlying, for example, photo- and chemotaxis, foraging and feeding, as well as learning and memory (Anreiter & Sokolowski, 2019; Eschbach & Zlatic, 2020; Louis, 2019; Miroschnikow et al., 2020; Thum & Gerber, 2019). Notably, focusing on larval food search strategies, Marla Sokolowski and co-workers were able to pinpoint for the first time the genetic and molecular bases for a naturally occurring behavioral polymorphism (Anreiter & Sokolowski, 2019; Osborne et al., 1997; Renger et al., 1999; Sokolowski, 1980). Furthermore, exploiting the natural transparency of the larval cuticle, Schroll et al. (2006) provided the first report on optogenetic manipulation of central brain neurons as a means of modulating behavior in freely moving animals. All these efforts benefit from an expanding repertoire of genetic tools for the combinatorial labeling and manipulation of subsets of neurons or single cells (Caygill & Brand, 2016; Simpson & Looger, 2018; Venken, Simpson, et al., 2011). With a complete atlas of the neurons in the larval brain and their chemical-synapse connectome within reach (Eichler et al., 2017; Gerhard et al., 2017; Li et al., 2014; Schneider-Mizell et al., 2016), recent studies have exploited the ease, completeness and precision with which larval neuronal structure-function relationships can be analyzed (Almeida-Carvalho et al., 2017; Eschbach et al., 2020; Eschbach & Zlatic, 2020; Jovanic et al., 2016; Miroschnikow et al., 2018; Ohyama et al., 2015; Saumweber et al., 2018; Thum & Gerber, 2019). These analyses are further facilitated by the development of a ‘standard brain’ to register patterns of transgene expression (Muenzing et al., 2018). In all these cases, the respective anatomical data were acquired from non-cleared, dissected brains based on reporter gene expression or immunofluorescence, or by electron microscopic serial sectioning and subsequent 3D reconstruction. However, the utility of these valuable data sets remains limited when it comes to relating them to behavior, simply because a corresponding account of the body, the ultimate behavioral actuator, is not available. Indeed, database documentations of Gal4-drivers specify their expression patterns only within the larval brain and ventral nerve cord (VNC) (this is the case for adult *Drosophila,* too: Jenett et al., 2012; Li et al., 2014; Muenzing et al., 2018). A routine procedure allowing for the systematic detection of transgenic reporter gene expression within the context of the whole body of larval *Drosophila* is therefore desirable.

Whole-body visualizations of labeled cells are established for embryonic *Drosophila* and have proved highly useful in developmental biology. Although by the end of embryogenesis the secreted cuticle strongly interferes with antibody labeling or *in situ* hybridization, transgenic reporters expressed within the body wall can still be detected during larval stages without the need for dissection (Aberle et al., 2002; Kakanj et al., 2020; Medina et al., 2006; Rasse et al., 2005; Zito et al., 1999). Due to tissue opacity, however, assessment of inner organs and substructures becomes increasingly inefficient with larval growth.

To overcome opacity, tissue clearing methods compatible with fluorescence labeling have been established, which, by equalizing differences in the scattering of light within tissues, generate transparency while largely preserving the structural and molecular integrity of the samples (Richardson & Lichtman, 2015, 2017). This has enabled the visualization of individual, fluorescently labeled cells within whole bodies and whole-organ samples from embryonic and adult mice (Belle et al., 2014; Cai et al., 2019; Chung et al., 2013; Susaki & Ueda, 2016). Combined with light sheet microscopy (Dodt et al., 2007) and pipelines for handling large data sets, these tissue clearing methods bear great potential for profiling normal and abnormal circuits and cellular arrangements in the mammalian brain and body (Ueda et al., 2020). However, only a few clearing-based protocols tailored to adult *Drosophila* are available (Jährling et al., 2010; Masselink et al., 2019; McGurk et al., 2007; Pende et al., 2018). Indeed, single-cell resolution in the adult brain has so far only been documented by Pende et al. (2018). While that study and Masselink et al. (2019) focused on adult *Drosophila* and other species, respectively, they also briefly report proof-of-principle applications of their protocols for *Drosophila* larvae. A detailed clearing protocol for the immunolabeling and detection of cells in the context of the whole larval body was presented by Manning and Doe (2017). This procedure, however, takes 12-16 days and requires considerable skill in handling, as well as expensive and toxic reagents. It is thus not ideal for routine use or screening approaches. In this context, we decided to develop a quick and user-friendly procedure for whole-body multi-fluorescence imaging of third-instar *Drosophila* larvae at cellular resolution.

## Materials and Methods

### Flies and fly husbandry

Flies were raised at 25°C, 50% relative humidity in a 12h/ 12h dark-light cycle and on food medium containing agar-agar (0.83% w/v), mashed raisins (4% w/v), baker’s yeast (6% w/v), semolina (5% w/v), sugar beet syrup (2.6%), honey (2.6% w/v) and nipagin (0.13%) (Erdmann et al., 2015). The following stocks were obtained from the Bloomington Drosophila Stock Center (BDSC): *y^1^ w*;; nsyb-Gal4* (stock #51635) (henceforth *nsyb-Gal4); w^1118^; vGlut^OK371^-Gal4* (#26160; Mahr & Aberle, 2006) (henceforth *OK371-Gal4); w^*^ dlg1^YC0005^* (#50859; Quinones-Coello et al., 2007) (as this is an EGFP-tagged protein-trap line henceforth *dlg1^EGFP-PT^); w*; UAS-mCherryCAAX* (#59021; Sens et al., 2010) (henceforth *UAS-mCherry-CAAX); w*;; UAS-mIFP-T2A-HO1* (#64181; Yu et al., 2015) (henceforth *UAS-mlFP-T2A-HO1); w*; IR76b-Gal4/CyO; TM2/TM6b* (#52610; prior to use, third-chromosomal balancers were outcrossed and the CyO balancer was replaced by a CyO-GFP balancer; Y. V. Zhang et al., 2013) (henceforth *IR76b-Gal4).* The Venus-tagged PMCA protein trap line *w^1118^;;; PMCA^CPTI001995^* was provided by the Kyoto Stock Center (#115256; Lowe et al., 2014) (henceforth *PMCA^Venus-PT^).* As a *Gal4* driver line specifically covering the APL neuron we used the previously described *ss01671-Gal4* (Saumweber et al., 2018) (henceforth *APL-Gal4).* The *w^1118^;; MB247:mCherry-CAAX* line was generated in the course of this study (see next section); the strain *y^1^ w^*^; MB247:mCherry-CAAX UAS-mIFP-T2A-HO1* was then generated by chromosomal crossover.

Larval progeny with genotypes of interest resulting from the crosses mentioned in the Results section were selected using a fluorescence stereomicroscope. For the starvation experiment shown in Figure S3, we kept the larvae on a wet filter paper at 70% humidity for 20 h prior to clearing. In all other cases, the larvae were non-starved.

### Generation of flies expressing mCherry-CAAX in the mushroom bodies

To generate flies expressing mCherry-CAAX in the mushroom bodies, i.e. under the control of the *MB247* promoter element (Schulz et al., 1996; Zars et al., 2000), we PCR-amplified the coding sequence for mCherry-CAAX from a single fly of the *UAS-mCherry-CAAX* stock (BDSC stock #59021; Sens et al., 2010) using the primers 5’-ctaaacaatctgcagGAATTCC-AAC ATGGTGAGC AAGGG-3’ and 5’-gctggaattaggcctTCTAGACTACATCAGGCAGCACTTCC-3’. The resulting PCR fragment was inserted by means of the cold fusion reaction (System Biosciences; CA, USA) into the EcoRI-XbaI-digested pCaSpeR-mb247-hs-GCamP3 vector (gift from T. Riemensperger, Cologne; courtesy of A. Fiala, Göttingen) to replace the GCamP3 sequence. The resulting construct was sequence-verified and used for germline transformation at BestGene Inc. (Chino Hills, CA, USA). One third-chromosome line was selected for further use (see previous section) upon validation of mushroom body expression.

### Clearing of third-instar larvae

Up to 30 third-instar larvae were collected using a brush, rinsed in tap water, and further treated in a glass block bowl (40 x 40 mm). The water was replaced by 4% bleach prepared from a fresh stock of 12% sodium hypochlorite (Roth, Karlsruhe, Germany, order no. 9062.1), diluted in tap water. After bleaching for 10 min at room temperature (RT), the larvae were washed 3 x 5 min in distilled water (dH_2_O). We note that bleaching is key for successful clearing and that aged sodium hypochlorite renders the cuticle insufficiently permeable whereas too high concentrations of sodium hypochlorite often lead to massive damage to the cuticle. Immediately after washing, dH_2_O was replaced by fresh 4% paraformaldehyde (PFA) in 0.1 M phosphate buffer (PB) at pH 7.4 for fixation. However, we increased pH to 9 to improve the detectability of EGFP-tagged Dlg (preliminary observations suggest that this might indeed be advantageous for other fluorescent proteins, too). The glass block bowl was covered with a lid and placed into a humid black box, and fixation continued with gentle shaking overnight at 4°C (3d protocol, Fig. S1C) or, alternatively, for 2 h at RT (1d protocol, not shown). Exposure to light was kept at a minimum for all the following steps. Fixed samples were briefly rinsed 3 x with 0.1 M PB containing 0.2% Triton-X-100 (PBT) and then washed 3 x 30 to 60 min in PBT. They were then either kept in PBT overnight at 4°C (3d protocol, Fig. S1C) until dehydration the following day, or were subjected right away to dehydration (1d protocol, not shown), which was achieved by a graded ethanol series. This series included 60-min incubations in 10% and 25% ethanol followed by 30-min incubations in 50%, 60%, 80% and 2 x 100% ethanol, with all ethanol dilutions adjusted to pH 9. The larvae were then optically cleared by replacing ethanol with ethyl cinnamate (ECi; Ethyl 3-phenyl-2-propenoate; Sigma-Aldrich, order no. 112372-100G). After 30-60 min of incubation, the ECi was replaced once with fresh ECi. The samples were from now on stored at RT in black boxes in a desiccator to avoid rehydration. After another 30 min the samples may appear fully transparent, but the clearing efficiency was found to increase with extended incubation time. Therefore, we recommend that the samples should not be subjected to imaging before an overnight incubation in ECi. Strikingly, and in line with a previous report (Klingberg et al., 2017), we found that cleared samples can be successfully imaged even after one year in ECi in a desiccator. We also note that in our experience, clearing may become less efficient, if >30 dehydrated larvae are subjected to ECi incubation in a single glass block bowl at a time. Finally, ECi-induced cuticular damage typically occurs, if at all, within the first 15 min after the addition of ECi. We therefore determined the rate of successfully processed, i.e. externally intact, larvae after this time point, simply by visual inspection on a stereomicroscope.

### Preparing phytagel blocks

Mounting of cleared samples onto a fully transparent substrate is a prerequisite for adequate imaging. To this end, ECi-cleared phytagel blocks were prepared. Phytagel powder (Sigma-Aldrich, P8169) was dissolved in tap water (dH_2_O is not suitable because cations are required for the cross-linking of the phytagel) to a final concentration of 0.75%, boiled for 1 to 2 min, and poured into a Petri dish to form a layer of approximately 1 to 1.5 cm thickness. The gel solidified within 20 min at RT. Blocks of approximately 1 x 1 x 1 cm were then cut out using a scalpel and transferred in sets of 5 to 7 to glass snap lid jars (25 ml). These blocks were dehydrated through an ethanol series in 2.5 h steps of 50%, 80% and 2 x 100% ethanol, which was then replaced by ECi. As in the case of the cleared larvae, the cleared phytagel blocks remain usable for at least several months when stored at RT in a desiccator.

### Mounting of larvae for light sheet microscopy

For light sheet microscopy (see next section), single cleared larvae were carefully transferred using forceps and placed ventral side up into a small furrow that had been cut with a scalpel into the surface of a phytagel block. The furrow may span the length of the phytagel block (ca. 1 cm) and should be 1-2 mm wide and deep, i.e. suitable to accommodate a larva at its full length. The loaded blocks were then transferred to a sample holder (microscope-specific fittings), which in turn was fixed on a mounting suspension (Fig. S1F). For imaging, this arrangement was clipped into a high-precision quartz glass cuvette filled with ECi, accessible via an opening within the microscope stage (Fig. S1G).

### Light sheet microscopy

Tiled image stacks were acquired on an UltraMicroscope II (Miltenyi Biotec, Bergisch-Gladbach, Germany) equipped with a Zyla 4.2 PLUS SCMOS camera (Oxford Instruments, Abingdon-on-Thames, UK). The setup was equipped either with an Olympus zoom body and a MVPLAPO 2x 0.5 NA dipping objective (Olympus, Tokyo, Japan) or an ultramicroscope tube for infinity-corrected objective lenses and a magnification changer, which allows for an additional 2x magnification. The dipping objectives for the tube were an LVMI-Fluor 4X/0.3 and an LVMI-Fluor 12X/0.53 (both from Miltenyi Biotec, Bergisch-Gladbach, Germany), or an HCX APO 20X/0.95 IMM (Leica Microsystems, Wetzlar, Germany). These objectives are suitable for imaging in ECi by matching its refractive index (RI) of 1.558. Furthermore, they provide a large working distance. Light from a supercontinuum EXW-12 extreme laser (NKT Photonics, Birkerød, Denmark) was guided through excitation filters (AHF Analysentechnik AG, Tübingen, Germany) and through triple sheet optics to illuminate samples from either just one side or from both sides. For data acquisition ImSpector software (version 5.0.285 – 7.0.124, Miltenyi Biotec, Bergisch-Gladbach, Germany) was used. Images acquired using two-side illumination were automatically fused with the built-in *blend* algorithm. In some cases, the built-in *dynamic focusing* function was used to achieve uniform z-resolution across the width (X-axis) of the sample. Resulting image parts were fused automatically through the built-in *contrast adaptive* algorithm. For all images presented in this study, acquisition parameters such as filters, objectives, z-slice numbers, z-step sizes, number of tiles, and exposure times are provided in Supplementary Table 1. With a format of 2048×2048 and the bit-depth set to 16-bit, the resulting data size was about 40-70 gigabytes per channel. For stitching, an overlap of tiles by 10% or 15% of the imaging format was used. Depending on the parameter settings, the time taken to image an entire larva varied considerably (Supplementary Table 1).

### Confocal laser scanning microscopy

For confocal laser scanning microscopy (CLSM), a phytagel block with a larva, prepared as described above, was transferred to a glass Petri dish to which it was glued with KWIK-SIL (World Precision Instruments, Berlin, Germany). The Petri dish was filled with ECi and then placed on the stage of an upright CFS 6000 microscope equipped with a TCS-SP5 AOBS scan head (Leica Microsystems, Mannheim, Germany). The scanning format was set to 2048×2048 and an optical zoom of 3 was applied. Imaging was performed with the same dipping objective HCX APO 20X/0.95 IMM (Leica Microsystems, Wetzlar, Germany) as was used for light sheet imaging. For details on imaging parameters see Supplementary Table 1.

### Post-imaging

The 16 bit ome.tif files from the light sheet microscope and the .lif-files from the CLSM were converted by the Imaris file converter (version 9.21-9.51, Bitplane, Zurich, Switzerland) into ims-files. Subsequently, single tiles were stitched using the Imaris stitcher software (version 9.21-9.51, Bitplane, Zurich, Switzerland), before being further processed and visualized. Inspection and rendering of data sets was done in Imaris (version 9.21-9.51, Bitplane, Zurich, Switzerland). To remove spurious, phytagel-derived signals from outside the larvae, an oversized surface was created in Imaris and the respective channel was masked with this surface to delete the signals. The *ortho slicer* tool in Imaris was used to restrict volumes in the z-direction to improve the representation of structures that were otherwise covered in the context of the whole body. All 3D images were generated in Imaris using the *snapshot* function. 2D maximum-intensity projections were generated in Fiji (Schindelin et al., 2012). Movies were produced in Imaris with the *key frame animation* tool, and Adobe Premiere Pro 2020 (Adobe Inc., San José, California, USA) was used for cutting and labeling. The still framing at the end of Movie 6 was produced in arivis Vision4D (version 3.4.0, arivis AG, Munich, Germany).

All processing of image data was performed on an HP Z840 workstation equipped with an Intel XEON CPU E5-2687Wv4, with 256 gigabyte RAM, the graphic board NVidia GTX 1080Ti, 2 PCI-SSD of 1 terabyte each for temporary folders and an 8 terabyte RAID10-SSD-Drive for storing the data. We recommend using a fast temporary folder on a separate SSD for smooth 3D operations.

## Results

### A user-friendly clearing procedure for third-instar *Drosophila* larvae

In a recent study we applied the organic solvent-based clearing protocol 3DISCO (Erturk et al., 2012) to visualize the nervous system in the context of the whole body of a third-instar larva (Saumweber et al., 2018). However, this method does not lend itself to routine use because i) the vast majority of larvae turned black during fixation in 4% paraformaldehyde (PFA), suggestive of melanization due to ruptured crystal cells; ii) a 3-day fixation is required; iii) it involves rather toxic agents (tetrahydrofuran, THF; dibenzylether, DBE); iv) fluorescence fades within a few days; and v) only about 1% of the larvae retain full integrity. The procedure that we introduce here overcomes these problems (Fig. S1; for a detailed protocol see the Materials and Methods section). As an initial step, a short bleach wash (Manning & Doe, 2017) was found to prevent blackening and allowed for a substantial shortening of the subsequent PFA fixation to a single overnight step at 4°C, or a 2h step at room temperature. We furthermore replaced THF and DBE by the less toxic clearing reagent ethyl-3-phenylprop-2-enoate (a.k.a. ethyl cinnamate, ECi) (Klingberg et al., 2017). Following thorough, ethanol-based dehydration, 2h incubation in ECi at room temperature resulted in satisfactory clearing of the larvae (Fig. S1; afterwards, the specimens can be stored using fresh ECi). As quantified for N= 25 runs with ≥ 30 larvae each, the average percentage of animals that endured the procedure without noticeable bodily damage was 19.84 ± 8,52%. Among the remaining larvae, there were always several specimens displaying only minor cracks or kinks such that they were still suitable for inspection. Depending on the selected conditions for fixation and post-fixation washes, the entire procedure takes between 1-3 days, with actual handling times of only about 2h (Fig. S1). We next assessed the compatibility of this protocol with imaging of various genetically encoded fluorescent markers. In accordance with our labs’ main research interests, we focused on the nervous system and paid particular attention to the visualization and integrity of nerve fibers. Unless mentioned otherwise, these assessments were carried out on a light sheet microscope.

### Detection of green and red fluorescent proteins in the whole-body context

To evaluate the detectability of proteins with an enhanced green fluorescent protein (EGFP) tag, we applied our protocol to larvae from a protein trap line in which the scaffold protein Dlg is expressed as an EGFP fusion protein in its endogenous pattern *(dlg1^EGFP-PT^,* Quinones-Coello et al., 2007). Hallmarks of Dlg expression in the periphery, most notably at NMJs, were detectable even without clearing (Fig S2A). However, elements and details of Dlg expression deeper inside the body of the larvae, for example in the neuropil of the central nervous system (CNS) and at septate junctions of the salivary glands, were revealed only after clearing (Fig. S2B-D) (Bachmann et al., 2010; Woods & Bryant, 1991). Along with distinct Dlg^EGFP^ signals, we noticed green fluorescence associated rather uniformly with most inner organs and tissues. This reflects autofluorescence (AF), as was also detectable in animals without EGFP expression (not shown). Although this is detrimental to the detection of weak EGFP signals, we reasoned that such green autofluorescence is actually useful in visualizing body structures as ‘landmarks’ in animals that transgenically express red fluorescent proteins as a target signal. As a proof of principle, we used the pan-neuronal driver *nsyb-Gal4* to express a membrane-tethered red fluorescent protein (mCherry-CAAX: Sens et al., 2010). Cleared *nsyb-Gal4>UAS-mCherry-CAAX* larvae showed strong mCherry-CAAX signals within the CNS and along peripheral nerve fibers (Fig. 1A), while red AF remained negligible with the exception of signals from food particles in the digestive tract (for more details see Fig. S3). In turn, green AF (Fig. 1B) can be used as a means of landmarking the target mCherry-CAAX signals, as evident from the enrichment of the mCherry-CAAX signal in the neuropil region of the larval CNS relative to the surrounding cell body cortex area in the green channel (Fig. 1C-C”).

**Figure 1:**
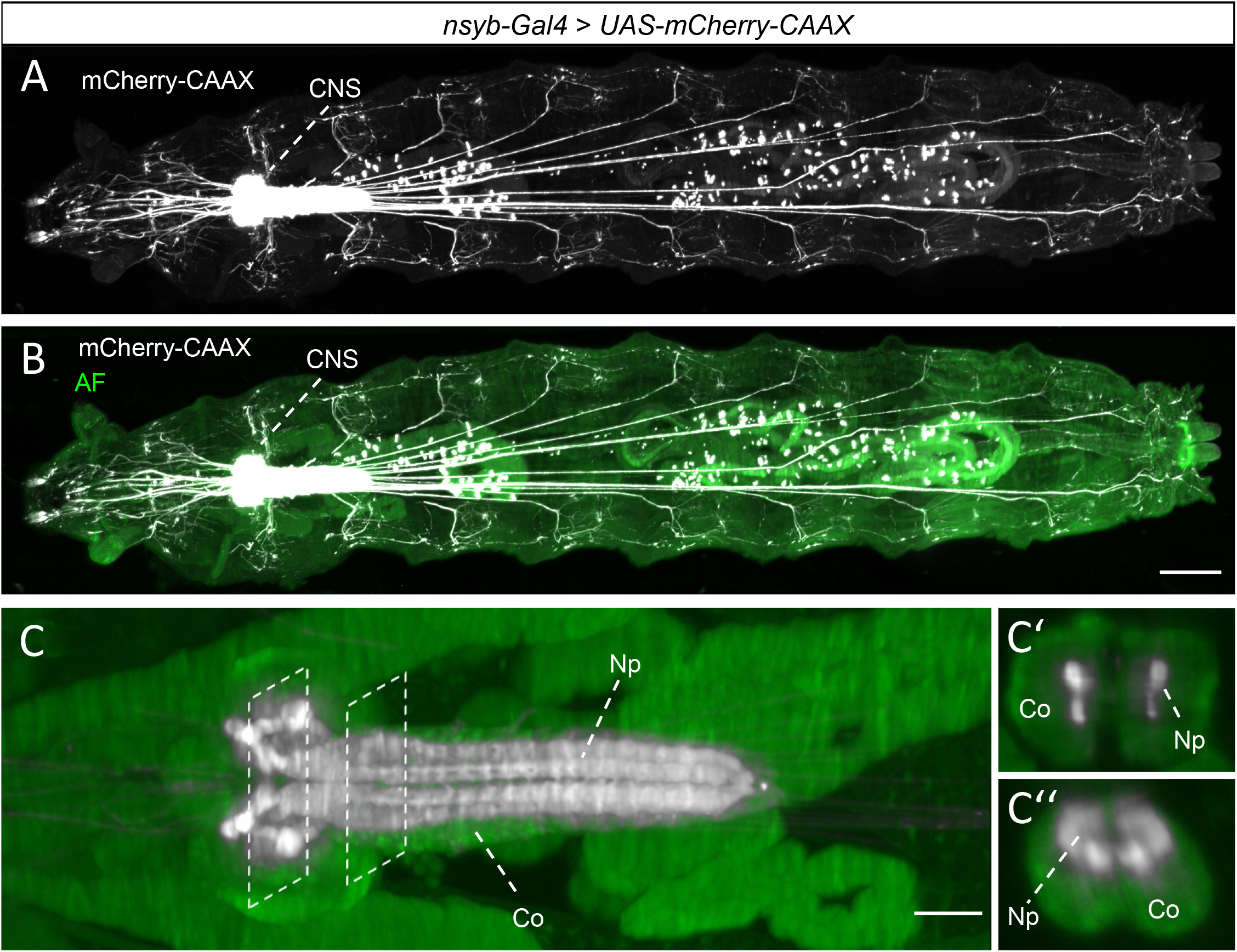
Detection of red fluorescent mCherry combined with green autofluorescence in the whole-body context. **(A)** Maximum intensity projection of an entire larva with pan-neuronally driven (nsyb-Gal4) mCherry-CAAX, acquired on a light sheet microscope with a 4x objective and an optical zoom of 2. Image contrast settings were adjusted to visualize nerve fibers in distal regions, resulting in the oversaturated appearance of the central nervous system (CNS). Top view, rostral to the left. **(B)** Same specimen as in (A) but with mCherry-CAAX labeling merged with green autofluorescence (AF). **(C-C‘‘)** Close-up of the CNS of the same specimen as in (A, B) in a volume restricted to 250 μm z-thickness. Image contrast settings were reduced as compared to (A, B), to show the enrichment of the mCherry-CAAX signal in the neuropil (Np) relative to the cell body cortex (Co). Stippled frames in (C) indicate positions of the cross sections in (C‘, C‘‘). Scale bars: 200 μm in (B), and 50 μm in (C). For this as well as for all other figures, further details on imaging parameters are given in Supplementary Table 1.

### Long-distance neuronal projections in the whole-body context

We next asked whether the present procedure leaves long-distance projections intact and traceable. This is essential for mapping out, within the context of the whole body, the connections between the CNS, the motor effectors and the sensory periphery, as well as within and between non-neuronal organ systems. We used the driver strain *IR76b-Gal4* to express mCherry-CAAX in external and pharyngeal larval chemosensory neurons and focused our attention on the fibers linking them to their target regions (the expression pattern of *IR76b-Gal4* in cephalic sensory neurons and their target regions in the CNS has itself been characterized in detail before: Croset et al., 2016; Rist & Thum, 2017). Indeed, intact fibers originating from sensory organs in the larval head could be traced all along their route to their target area within the CNS, the subesophageal zone (SEZ) (Fig. 2A, A’). Moreover, we noticed discrete neurites ascending from segmentally recurring somata located in the ventral periphery of the larval body (Fig. 2A’’). In fact, *IR76b-Gal4* has recently been shown to express in a subset of ventral tracheal dendritic neurons (v’td1) that serve the perception of interoceptive signals (Qian et al., 2018). Movie 1 traces these v’td1 axonal projections from the periphery to the VNC, indicating that even these filigree structures have withstood the clearing procedure. Likewise, we observed that in cleared *nsyb-Gal4>UAS-mCherry-CAAX* larvae, motor neuron projections to the hindgut (W. Zhang et al., 2014) remained traceable (Fig. 2B-B’’ and Movie 2).

**Figure 2:**
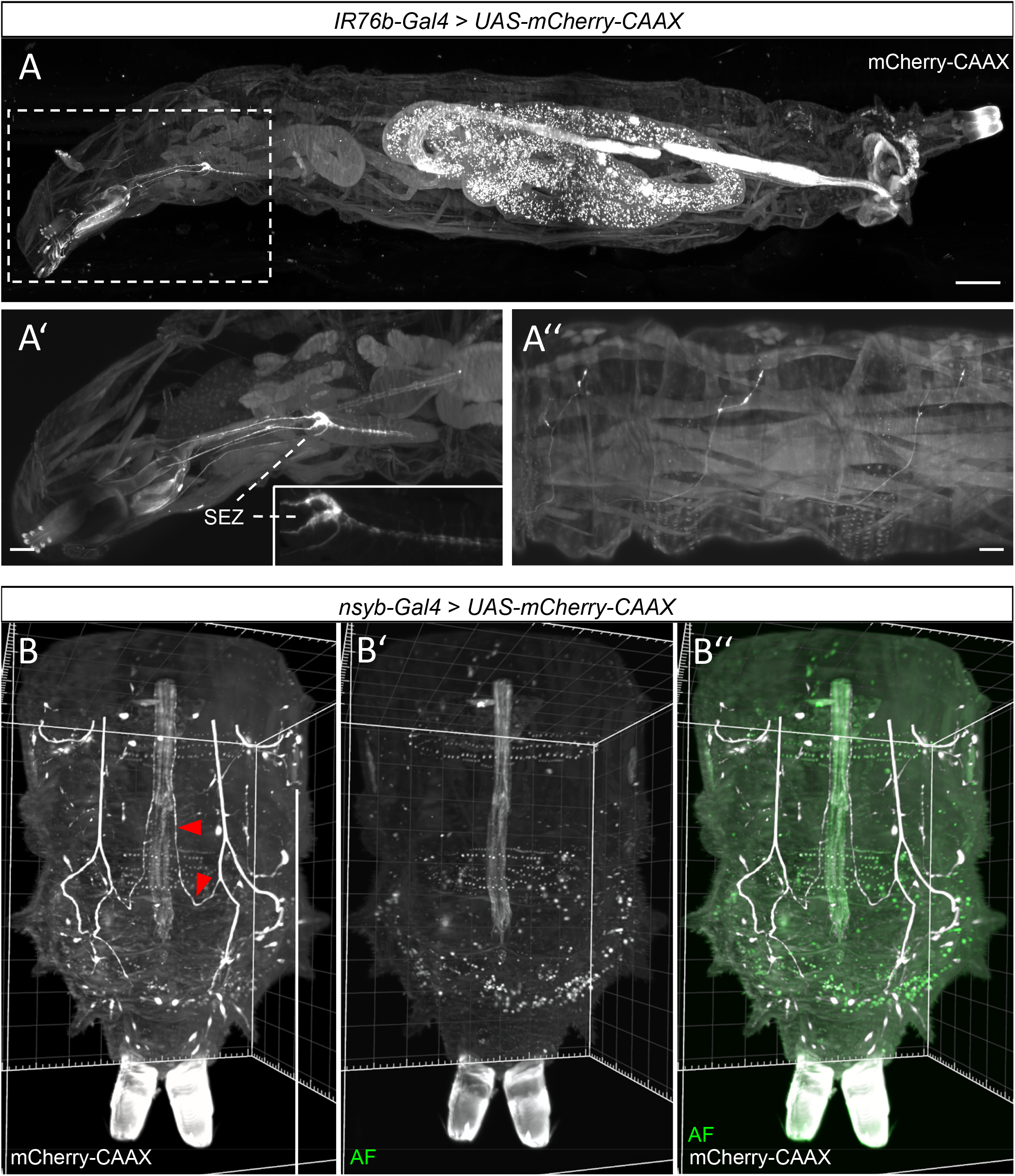
Long-distance neuronal projections in the whole-body context. **(A)** Maximum intensity projection of an entire larva with peripherally driven (IR76b-Gal4) mCherry-CAAX, acquired on a light sheet microscope with a 2x objective and an optical zoom of 4. The area indicated by the stippled frame in (A) is shown as a close-up in **(A‘)**, to highlight the traceability of sensory projections from the head region to the subesophageal zone (SEZ) in the CNS. The inset in (A’) shows a further magnification of the SEZ. **(A“)** shows the labeling of v’td1 axonal projections within three abdominal segments of the same specimen. (A, A‘) represent side views, rostral to the left. (A“) represents a ventro-lateral view, again rostral to the left. (A‘, A“) are restricted volumes (substacks). See also Movie 1. **(B-B‘‘)** Caudal region of a larva with pan-neuronally driven (nsyb-Gal4) mCherry-CAAX, showing motor neuron projections innervating the hindgut (arrowheads). Autofluorescence in green: AF. Caudal is to the bottom. See also Movie 2. Scale bars: 200 μm in (A) and 100 μm in (A‘, A‘). Grid spacing in (B): 100 μm.

### Monomeric IFP as a superior, clearing-resistant reporter

To further evaluate the present protocol for its versatility in multicolor fluorescence imaging, we turned to the monomeric infrared fluorescent protein (mIFP). Specifically, we used the recently developed reporter strain *UAS-mIFP-T2A-HO1*, which allows for Gal4-driven coexpression of mIFP and hemoxygenase HO1, an enzyme that converts endogenous heme to biliverdin, which in turn serves as a chromophore required for mIFP fluorescence (Yu et al., 2015). As illustrated for pan-neuronally expressed mIFP in Figure 3 and Movie 3, we found this reporter to be well detectable in cleared larvae and even to outperform mCherry-CAAX in terms of signal-to-noise ratio. However, Gal4-independent fluorescence in the gastrointestinal tract was also observed in the near-infrared range (Fig. 3A). As these signals are highly variable between individuals in terms of their intensity and location (not shown), we reasoned that they might originate from food particles. Indeed, starvation prior to clearing substantially reduces these signals (Fig. S3). As we did not notice a substantial increase in bodily damage in starved larvae, starvation protocols varying in duration or variations in diet may be considered as ways of modifying the procedure.

**Figure 3:**
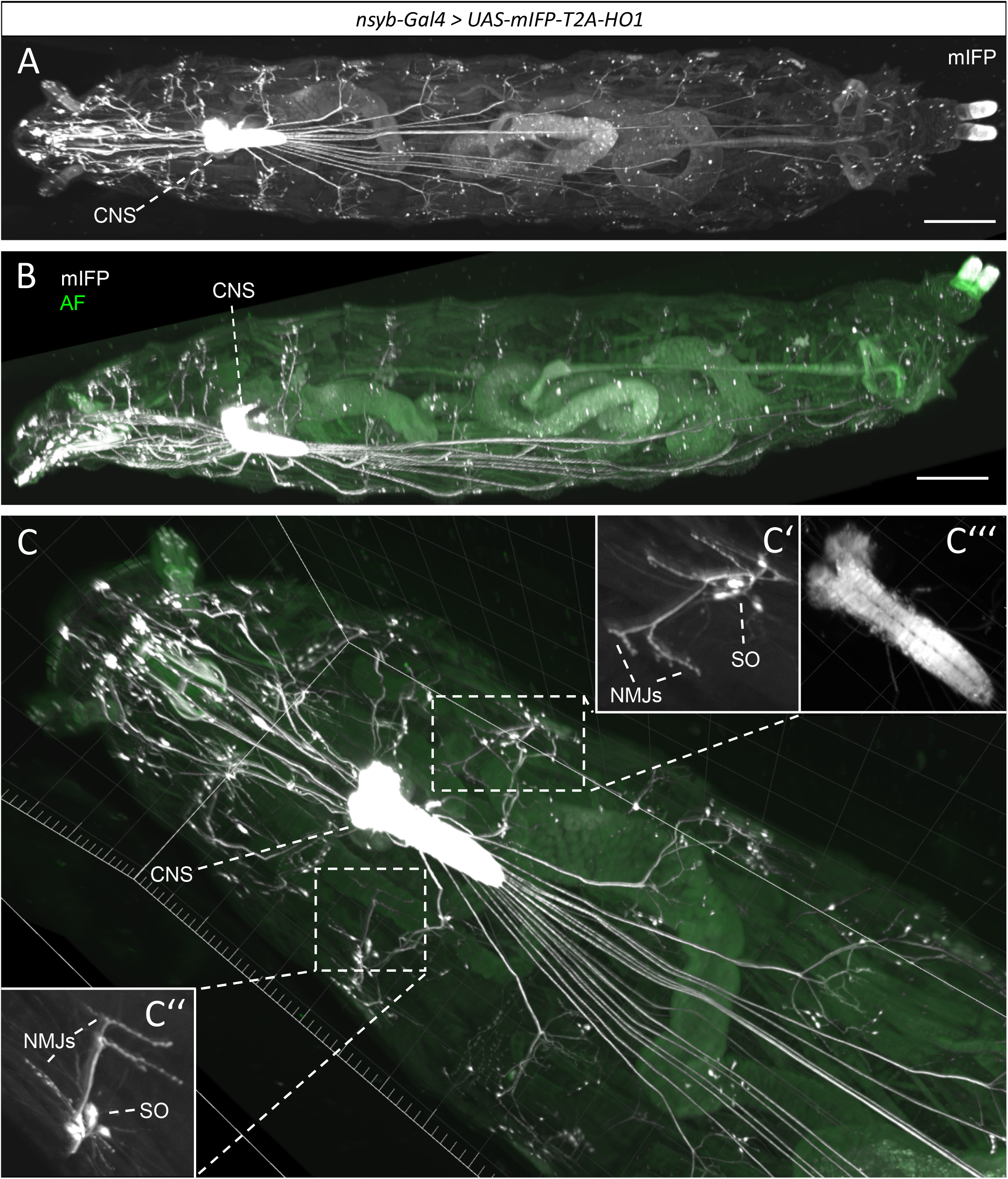
Monomeric IFP as a superior clearing-resistant reporter. **(A)** Three-dimensional rendering of an entire larva with pan-neuronally driven (nsyb-Gal4) mIFP, acquired on a light sheet microscope with a 12x objective. Image contrast settings were adjusted to visualize nerve fibers in distal regions, resulting in the oversaturated appearance of the central nervous system (CNS). Note the low level of autofluorescence in the near-infrared channel. Top view, rostral to the left. **(B)** The same specimen as in (A) but in a side view and with the mIFP signal merged into the autofluorescence signal in the green range for better landmarking. **(C-C‘‘‘)** Same specimen as in (A, B), showing a close-up of the rostral region including head, thorax and the first three abdominal segments, in a top view with rostral towards the upper left. (C‘, C‘‘) show neuromuscular junctions (NMJs) next to cell bodies of sensory organs (SO). (C‘‘‘) shows a contrast-adjusted image of the CNS. Scale bar: 400 μm in (A and B). Grid spacing in (C): 200 μm. See also Movie 3.

### Two-color fluorescence imaging in the whole-body context: visualizing connected neurons

The suitability of both mCherry-CAAX and mIFP in the present protocol allowed us to express these two reporters in distinct sets of neurons. To exploit this possibility for two sets of neurons known to be synaptically connected, we generated transgenic flies expressing mCherry-CAAX under the control of the mushroom-body-specific promoter element MB247 *(MB247:mCherry-CAAX)* and recombined them with the *UAS-mIFP-T2A-HO1* reporter; flies from this strain were then crossed to *APL-Gal4,* which drives expression specifically in the GABAergic anterior paired lateral (APL) neurons (Saumweber et al., 2018). APL is known to be synaptically and functionally connected with the mushroom body intrinsic neurons (Eichler et al., 2017; Masuda-Nakagawa et al., 2014; Saumweber et al., 2018). As expected, mCherry-CAAX was highly expressed in the mushroom bodies (Fig. 4A, B), with a striking enrichment in the neuropil and less pronounced, yet discernible expression in the cell bodies (Fig. 4B). In turn, the mIFP signals revealed the known anatomical features of APL, including a very large cell body, a thick and characteristically bent primary neurite, and projections onto both the mushroom body calyx, i.e. the dendritic compartment of the mushroom body intrinsic neurons, and select compartments of the mushroom body lobes (Fig. 4B’, B’’ and Movie 4; a second example of two-color fluorescence imaging, using *IR76b-Gal4* as the driver in neurons known not to be connected to the mushroom body, is shown in Movie 5). Notably, consecutive scanning of one and the same specimen on a light sheet and then on a confocal laser scanning microscope (CLSM) (Fig. 4C-C’’) revealed that the present procedure is suitable for confocal imaging, too.

**Figure 4:**
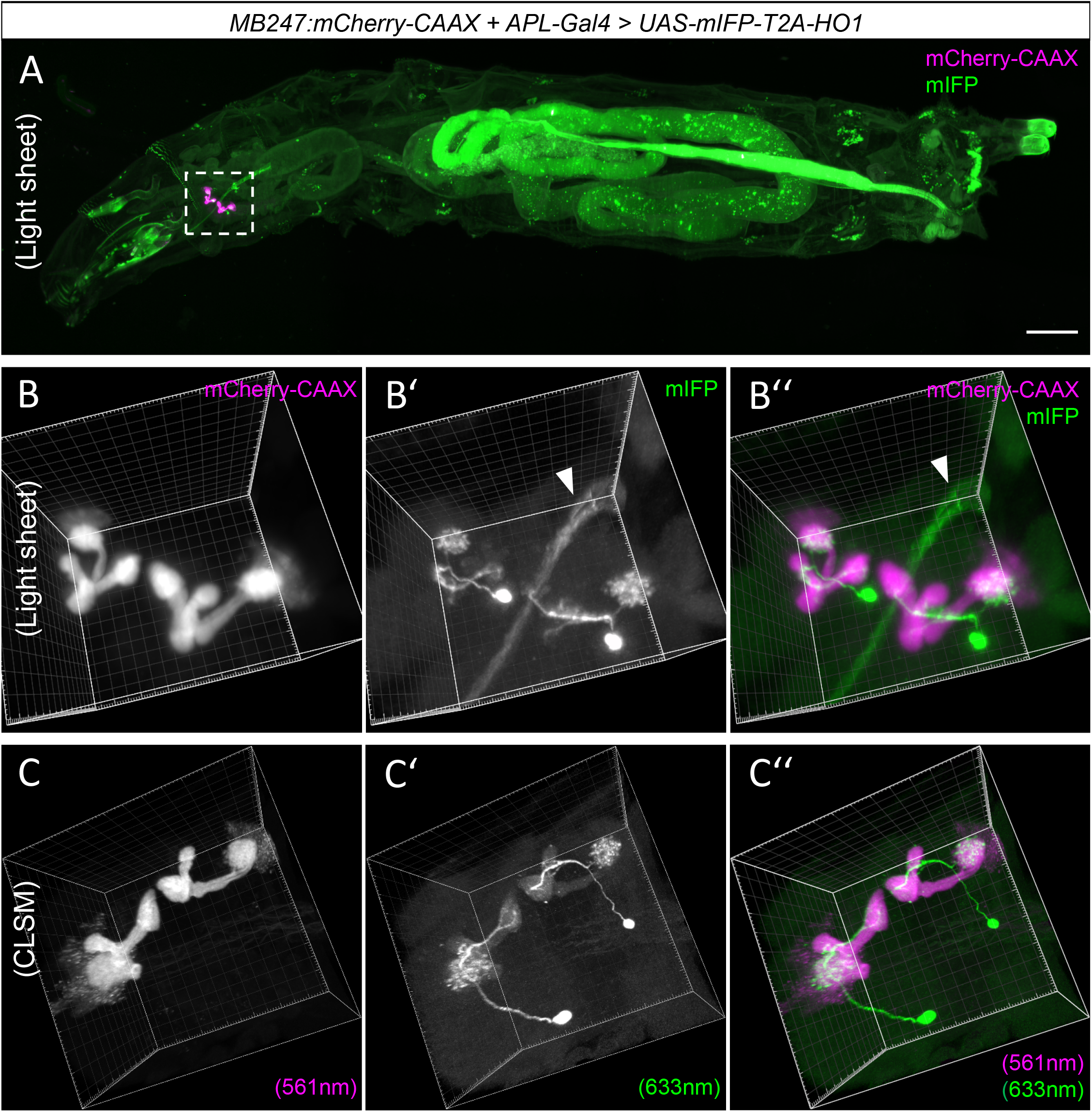
Two-color fluorescence imaging in the whole-body context: visualizing connected neurons. **(A)** Maximum intensity projection of an entire larva expressing mCherry-CAAX in the mushroom bodies (magenta; by means of the MB247 promotor) and mIFP in the APL neuron (green; by APL-Gal4), acquired on a light sheet microscope using a 2x objective and an optical zoom of 4. Top view, rostral to the left. Note that in this case high contrast settings for the mIFP channel were used to better visualize the APL neurons next to the bright mCherryCAAX signal within the boxed area. This results in an overrepresentation of the otherwise low autofluorescence in the mIFP channel in the gut (compare to Fig. 3A). **(B-B‘‘)** Close-ups of the volume indicated by the stippled frame in (A). Labeling in the pharynx (arrowheads in B‘, B‘‘) is due to autofluorescence. **(C-C‘‘)** Same specimen as in (B-B”), acquired on a confocal laser scanning microscope (CLSM), using a 20x objective appropriate for organic solvent-based media, shown at a different angle. Note that the use of a 633 nm laser on the CLSM required high laser power to achieve reasonable emission from mIFP (excitation maximum 680 nm), which resulted in weak cross talk between the channels such that a faint labeling of the mushroom bodies is visible in (C‘). Numbers in brackets indicate excitation wavelength. Scale bar in (A): 200 μm. Grid spacing in (B-C”): 10 μm. See also Movies 4, 5.

### Three-color fluorescence imaging in the whole-body context

A yellow fluorescent variant of GFP, called Venus, was recently used to establish a large number of protein trap lines (Lowe et al., 2014). We applied the present protocol to larvae from one such strain, *PMCA^Venus-PT^,* which expresses a Venus-tagged plasma membrane calcium ATPase (PMCA) from the endogenous *PMCA* locus. To achieve triple labeling, *PMCA^Venus-PT^* flies carrying either the Gal4 driver *nsyb-Gal4* or *OK371-Gal4* were crossed to flies carrying both *MB247:mCherry-CAAX* and *UAS-mIFP-T2A-HO1.* We found that the Venus-tagged PMCA was clearly detectable in the nervous system of larval progeny (Fig. 5A, B) and, importantly, that the Venus signal remained clearly separable from both the mCherry-CAAX signal in the mushroom bodies (Fig. 5A’, 5B’) and from the mIFP signal established either pan-neuronally (Fig. 5A” and Movie 6) or in motor neurons (Fig. 5B”, B’”).

**Figure 5:**
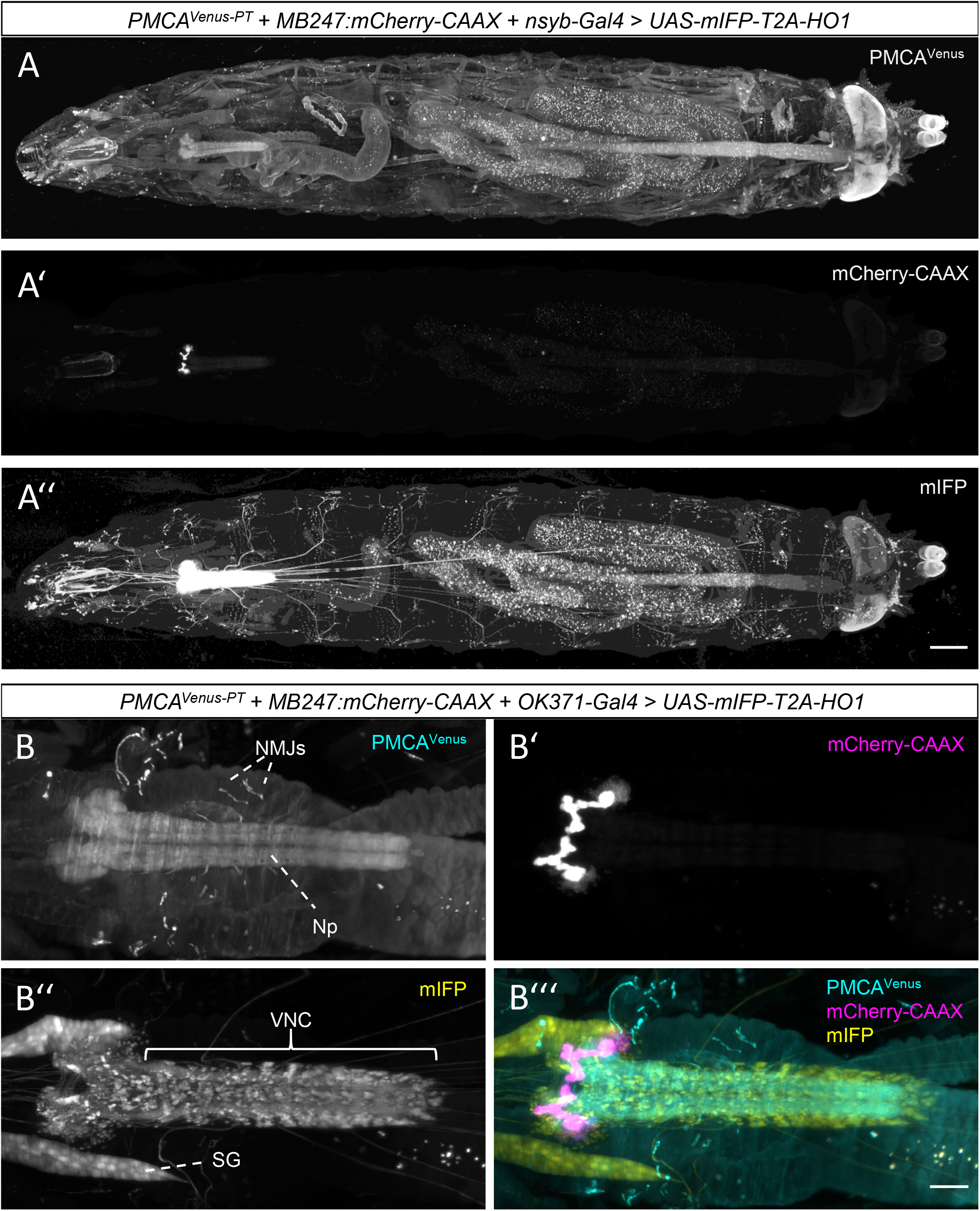
Three-color fluorescence imaging in the whole-body context. **(A-A’’)** Maximum intensity projection of an entire larva expressing a Venus-tagged PMCA protein from the endogenous PMCA locus (A), mCherry-CAAX from the MB247 promoter (A’), and mIFP driven by nsyb-Gal4 (A”); the data were acquired on a light sheet microscope using a 12x objective. Top view, rostral to the left. See also Movie 6. **(B-B’’’)** The central nervous system of a larva expressing a Venus-tagged PMCA from the endogenous PMCA locus (B; note the enrichment of the signal in the neuropil (NP) and on neuromuscular junctions (NMJs)), mCherry-CAAX from the MB247 promoter (B’), mIFP as expressed in motor neurons by OK371-Gal4 (B’’; note the individual cell bodies in the ventral nerve cord (VNC) and expression in the salivary glands (SG)), as well as the three-color merge of these signals (B’’’), acquired with a 20x objective (PMCAVenus: turquoise; mCherry-CAAX: magenta; mIFP: yellow). Top view, rostral to the left. Scale bars: 200 μm in A’’ and 50 μm in B’’’.

## Discussion

Pioneered by Werner Spalteholz a century ago, tissue clearing proved to be a groundbreaking technique in microscopic anatomy (Eisenstein, 2018; Spalteholz, 1911) once optimized protocols were combined with state-of-the-art cell labeling, microscopy, and image data processing (Ueda et al., 2020). This paved the way for striking insights into the morphology and cytoarchitecture of relatively large, intact, and complete organs, including mammalian brains and indeed their neuronal projections throughout the body (Cai et al., 2019). Given that these approaches work for animals the size of a mouse and ‘through skin and bone’, it might have seemed trivial to apply them to much smaller animals such as *Drosophila,* too. The cuticular exoskeleton of arthropods, however, proves to be a tricky obstacle to clearing. A recent protocol (Pende et al., 2018) has addressed this problem with remarkable success and was documented and optimized for pupal and adult stages.

In the present study we have developed a clearing protocol tailored specifically to third-instar larvae, user friendliness, and routine use. Important features of the present procedure include a brief initial incubation with fresh bleach, making the enzymatic digestion of the exoskeleton unnecessary (Masselink et al., 2019; Pende et al., 2018), and the use of ECi (Klingberg et al., 2017) instead of more toxic agents such as DBE or THF (Erturk et al., 2012) or hydrophilic reagents (Ueda et al., 2020; Gómez-Gaviro et al., 2020). ECi is not only a budget compound and approved as non-toxic. As an organic agent, it is particularly suitable for clearing lipid-rich tissues such as the larval fat body, and in this and other studies it has proven to be convenient in handling and fast-acting (Klingberg et al., 2017; Masselink et al., 2019; Huang et al., 2019). In agreement with Masselink et al. (2019), indeed, we found that clearing by ECi can be achieved within a few hours. In contrast, the clearing protocol by Pende et al. (2018), modified from the widely used CUBIC method, requires consecutive immersion of fixed larvae in two hydrophilic clearing solutions for 5-6 days. However, the latter method might be favored if no microscope objective suitable for imaging at an RI near that of ECi (1.56) is available. We further found that the ECi-cleared larvae retain both tissue- and fluorescence stability for at least one year.

The present procedure can be carried out within a day and requires minimal hands-on time. Moreover, the larvae can be processed in groups of up to 30 at a success rate of about 20% and mostly preserving long-distance neuronal projections. Damage often occurred at the step where ethanol was replaced by ECi, suggesting that efforts for further improvements might focus on this or the preceding steps. Although in our hands embedding the larvae into phytagel before ethanol dehydration did not lead to obvious improvements in success rate, the use of other alcohols such as 1-propanol, or of PBS to dilute alcohol during dehydration (Masselink et al., 2019), or a stepwise switch from alcohol to ECi may be considered. We would like to stress, however, that rather than the 20% success rate it is the image acquisition process that we consider to be limiting for large-scale screening approaches.

We placed particular emphasis on evaluating the method for its compatibility with microscopic inspection, not least by testing various fluorescent reporters. The widely used EGFP and its derivative EYFP/Venus work well, provided that expression levels are high enough to outshine autofluorescence, which is prominent in the respective range of wavelengths. Green autofluorescence, in turn, can be used to provide a tissue map for the projection of red and near-infrared fluorescence target signals from other reporters. Low-level reporter expression is indeed often encountered in EGFP- or Venus-based protein trap lines. Since our protocol is not compatible with signal enhancement by antibody labeling, we would like to mention available strategies for genomically replacing EGFP by other fluorescent proteins or transcriptional activators to increase the signal-to-noise ratio and thus to assess gene expression patterns more accurately (Diao et al., 2015; Venken, Schulze, et al., 2011). While our attempts to detect Gal4-driven DsRed after clearing remained unsatisfactory (not shown), we found that membrane-tethered mCherry-CAAX works particularly well as a reporter in cleared larvae. Furthermore, we found that soluble mIFP, which was introduced for live imaging (Yu et al., 2015), is considerably quenched in conventionally fixed samples (not shown). Our observation that it works well with our protocol thus came as a surprise and may prompt further trying out of additional, in particular red-shifted, reporters.

Our procedure allows the monitoring of various reporters that are expressed independently of each other in the same animal, and throughout the whole body. Here, we used the Gal4/UAS system for the targeted expression of one fluorescent reporter along with one or two other fluorescent proteins expressed under direct control of *cis*-regulatory elements *(PMCA^Venus-PT^, MB247:mCherry-CAAX).* The Gal4/UAS-system, however, could just as well be used in combination with the Q- and lexA/lexAop transactivator systems to exploit the combinatorial potential of the multitude of available driver lines for each of these systems (Venken, Simpson, et al., 2011).

The whole-body inspection of the expression patterns permitted by our protocol might do more than just enable a more comprehensive characterization of the larval anatomy. Rather, such whole-body assessment could have implications for functional analyses by means of, for example, RNAi or optogenetic effectors, because driver lines with well-characterized expression in any one organ, such as the brain, may also drive expression in cells or tissues elsewhere in the larval body. If such expression remains unrecognized, interpretation of phenotypes may unwittingly go astray. The present clearing protocol, we hope, will help to avoid such pitfalls. Particularly when combined with the relatively fast imaging on a light sheet microscope, this should be possible on a routine basis. While the equipment used for the present study allowed us to visualize neurons at cellular resolution, higher resolution can be achieved through thinner light sheets (Pende et al., 2018). Moreover, newly developed fluorescent proteins (Matlashov et al., 2020), especially those emitting in the far-red range, may well complement the present two- and three-color approaches to multi-fluorescent imaging.

As it stands, the present clearing procedure and imaging pipeline is a step towards contextualizing neurons and other cell types within the whole larval body. We note that the 3D representations as shown in our movies can be developed further towards virtual reality visualization. In combination with the standard brain developed by Muenzing et al. (2018) and integration with connectomic datasets (Eichler et al., 2017; Gerhard et al., 2017; Li et al., 2014; Schneider-Mizell et al., 2016), this should provide a powerful toolset for further research into the larva – and a study case into what a multimodal imaging approach to structure can do for functional analyses.

## Acknowledgements

Discussions with and comments from Eike Budinger, Nino Mancini, Torsten Stöter, Naoko Toshima, Werner Zuschratter (LIN), Hans Ulrich Dodt and Marco Pende (TU Wien), and expert technical assistance by Bettina Kracht (LIN) are gratefully acknowledged. We thank Rupert D.V. Glasgow (Zaragoza, Spain) for language editing and appreciate the supply of stocks by the Bloomington Drosophila Stock Center (Bloomington, Indiana, USA) (NIH P40OD018537) and the Kyoto Stock Center (Kyoto, Japan), and the supply of plasmid DNA by Thomas Riemensperger (University of Cologne).

