## Supplementary Figures S1-S3 for "A quick and versatile protocol for the 3D visualization of transgene expression across the whole body of larval *Drosophila*"

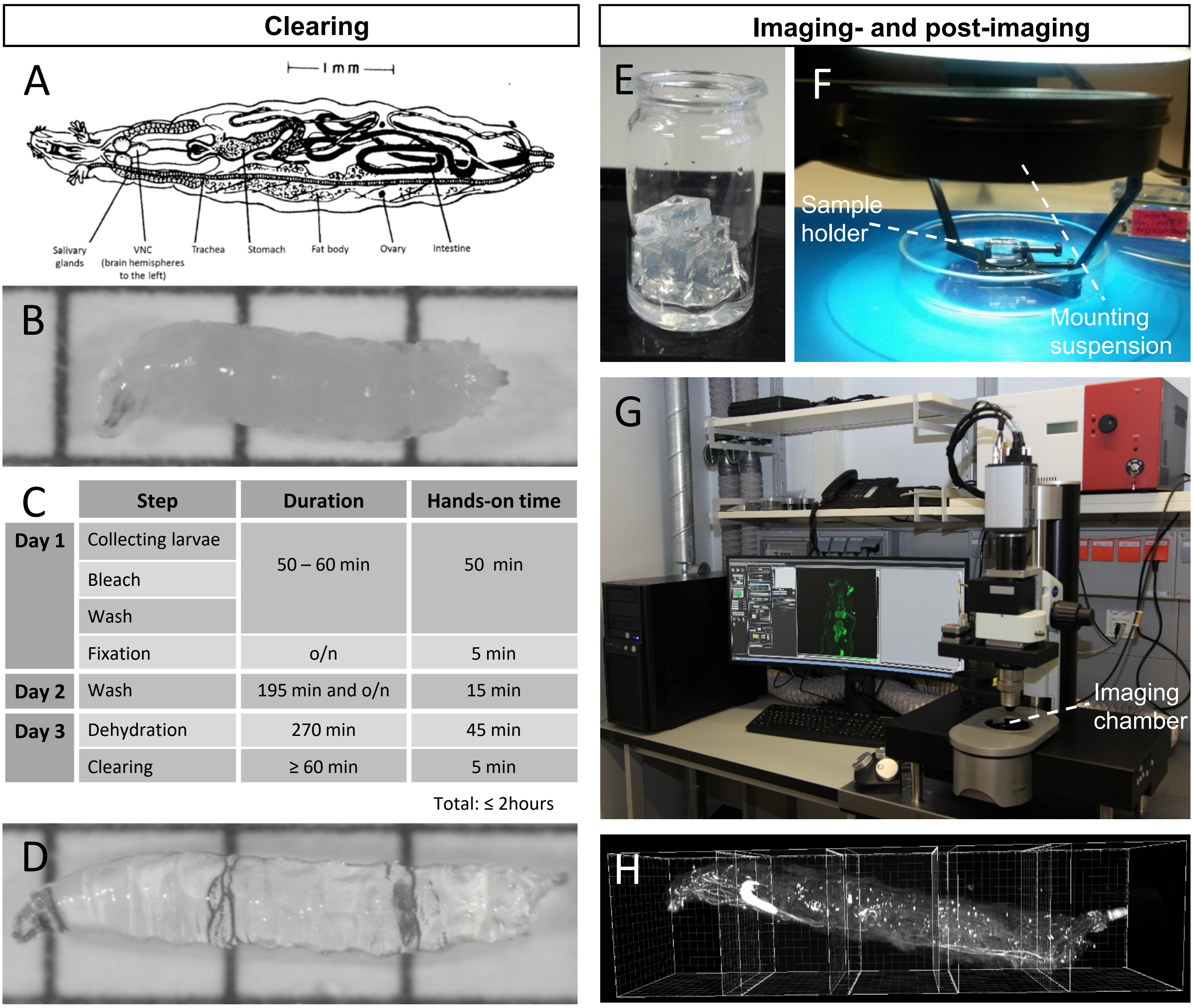


| | Step | Duration | Hands-on time |
|---|---|---|---|
| Day 1 | Collecting larvae | 50 – 60 min | 50 min |
| | Bleach | | |
| | Wash | | |
| | Fixation | o/n | 5 min |
| Day 2 | Wash | 195 min and o/n | 15 min |
| Day 3 | Dehydration | 270 min | 45 min |
| | Clearing | ≥ 60 min | 5 min |
| | | | Total: ≤ 2hours |

**Figure S1: A user-friendly clearing procedure for third-instar Drosophila larvae.**
**(A)** Schematic overview of anatomical relationships within the body of third-instar Drosophila larvae (from Demerec, M. & Kaufmann, B. P., Drosophila Guide: Introduction to the Genetics and Cytology of Drosophila melanogaster. Carnegie Institution of Washington, Washington, D.C., 1940); VNC refers to the ventral nerve cord. Note that tissues such as the peripheral nervous system, muscles and imaginal discs are omitted. **(B-D)** Clearing of third-instar larvae. (B) Larva on a 1.25 mm grid, prior to clearing. (C) Steps and time budget of the clearing procedure (3-day version), using ethyl cinnamate (ECi) as clearing solvent. o/n: overnight. (D) Larva after clearing. **(E-H)** Imaging and post-imaging for cleared larvae. (E) Examples of cleared phytagel blocks. (F) Larvae are embedded by placing them into a small slit on top of cleared phytagel blocks, which are then transferred to a sample holder for imaging on a light sheet microscope (G). (H) Due to the restricted field of view depending on the magnification of the objective used, acquired tiles (stacks) have to be stitched together during post-imaging, using a high-power workstation prior to 3D rendering. Tiles are indicated by so-called bounding boxes. A detailed description of the clearing, imaging and post-imaging is given in the Materials and Methods section.

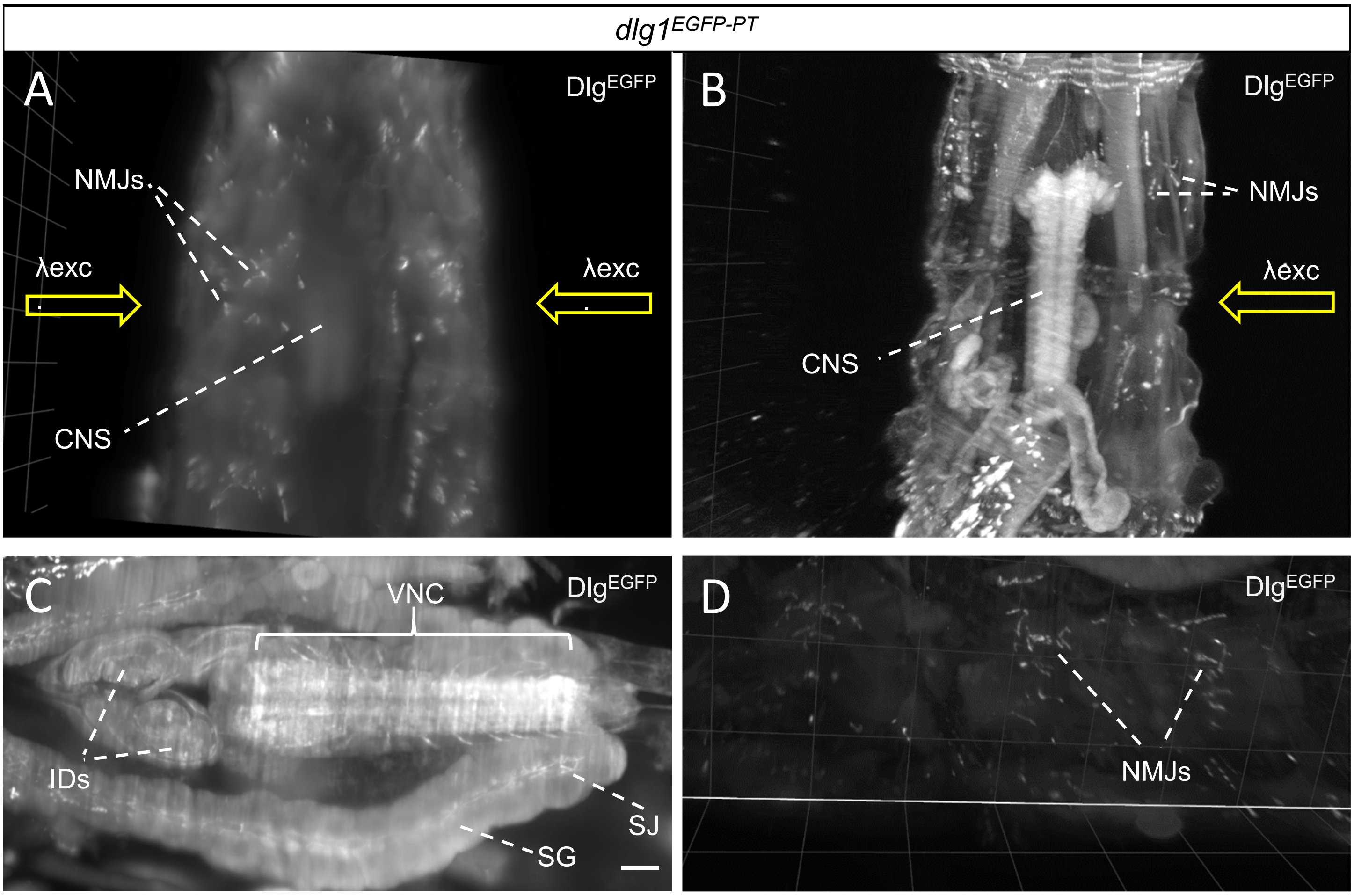


**Figure S2: Detection of green fluorescent proteins.**
**(A-D)** Green fluorescence from EGFP-tagged Dlg, expressed from the endogenous dlg locus in the protein-trap allele dlg1EGFP-PT as acquired on a light sheet microscope. A comparison is shown between a non-cleared larva (A) and a sibling ECi-cleared larva (B), displaying their thoracic segment T3 and abdominal segments A1 and A2, ventral view. Along with the larval samples, the phytagel blocks for embedding were non-cleared and ECi-cleared, respectively. For monitoring the non-cleared specimen under water, the objective was adjusted to an RI of 1.33 by mounting an appropriate dipping cap. Bidirectional excitation (arrows) is required for the non-cleared sample to visualize the neuromuscular junctions (NMJs) on both sides. The central nervous system (CNS) can hardly be discerned (A), whereas unilateral excitation is sufficient for the cleared larva to allow a more detailed view of EGFP-tagged Dlg not only at the CNS and NMJs (B), but also at imaginal discs (IDs), the ventral nerve cord (VNC), and the salivary glands (SG), including their septate junctions (SJ) (C, D). (C) shows a volume restricted to 250 µm z-thickness (substack) whereas in (D) the full volume is shown. Data were acquired with a 12x objective. Rostral up in (A, B) and to the left in (C, D). Scale bar in (C): 50 µm. Grid spacing is 100 µm in (A, B) and 200 µm in (D).

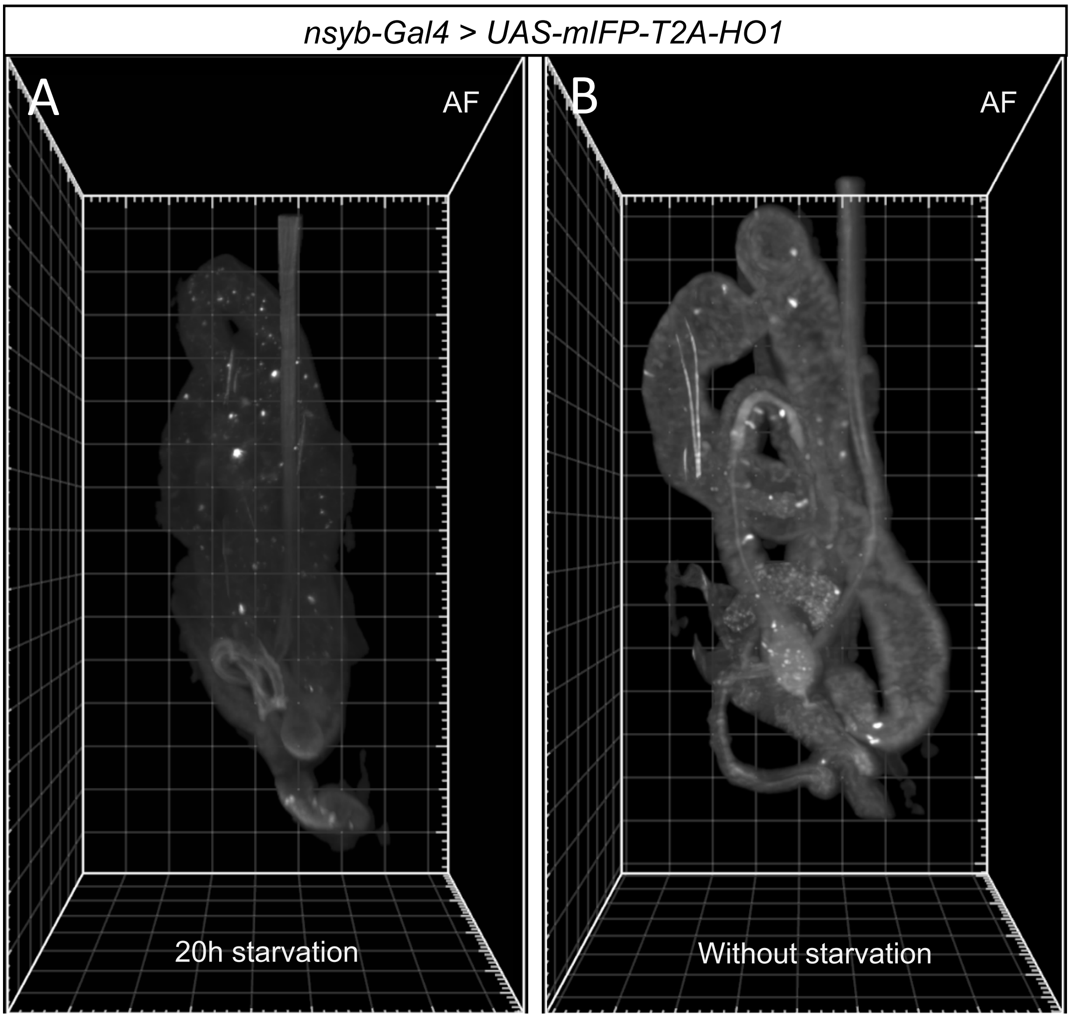


***Figure S3: Starvation reduces near-infrared fluorescence in the gut.***

**(A, B)** Comparison of near-infrared autofluorescence (AF) in the gut of larvae after 20h starvation (A) and without starvation (B), respectively. In these larvae, mIFP was expressed in neurons only (nsyb-Gal4). Fluorescence signals are substantially reduced upon starvation, suggesting that they originate from food particles. Data were acquired on a light sheet microscope using a 2x objective with an optical zoom of 2.5. Caudal to the top. Grid spacing: 100 µm.
