## Supplementary Table 1 for "A quick and versatile protocol for the 3D visualization of transgene expression across the whole body of larval *Drosophila*"

**Supplementary Table 1: Imaging parameters**

Abbreviations: exc.: excitation; em.: emission; corr.: corrected; dyn.: dynamic; illumin..: illumination; exp.: exposure; l/s: lines per second

| Figure | Fluorescence filter [nm] | | | | | | | | | | | Microscopy parameters | | | | | | | | |
|---|---|---|---|---|---|---|---|---|---|---|---|---|---|---|---|---|---|---|---|---|
| | Autofluorescence | | EGFP / Venus | | mCherry-CAAX | | mIFP | | Objective | | | | | | | | | | | |
| | exc. | em. | exc. | em. | exc. | em. | exc. | em. | Olympus zoombody | Infinity corr. tube | Leica CLSM | Optical zoom | z-step [µm] | z # | tiles # | dyn. focus | Illumin. sides | Exp. time [ms] | Acquisition time [min] | Data size [GB] |
| 1 A-C’’ | 470/30 | 520/40 | / | / | 570/20 | 625/30 | / | / | / | 4x | / | 2 | 1 | 854 | 3 | yes | 1 | 255 | 185 | 40 |
| 2 A-A’’ | / | / | / | / | 595/20 | 625/30 | / | / | 2x | / | / | 4 | 1 | 1191 | 4 | no | 1 | 200 | 26 | 37.2 |
| 2 B-B’’ | 681/24 | 720/24 | / | / | 595/20 | 625/30 | / | / | / | 12x | / | 1 | 1 | 872 | 1 | no | 1 | 162 | 8 | 13.6 |
| 3 A-C’’’ | 530/30 | 585/40 | / | / | / | / | 681/24 | 720/24 | / | 12x | / | 1 | 1 | 1537 | 6 | yes | 1 | 215 | 528 | 144 |
| 4 A-B’’ | / | / | / | / | 595/20 | 625/30 | 681/24 | 720/24 | 2x | / | / | 4 | 2 | 627 | 4 | no | 1 | 300 | 34 | 39.2 |
| 4 C-C’’ | / | / | / | / | 561 | 580-620 | 633 | 680-730 | / | / | 20x | 3 | 0.5 | 366 | 1 | / | / | 400 l/s | 96 | 6.1 |
| 5 A-A’’ | / | / | 500/20 | 535/30 | 595/20 | 625/30 | 681/24 | 720/24 | / | 12x | / | 1 | 1 | 1361 | 5 | yes | 2 | 199 | 1113 | 207 |
| 5 B-B’’ | / | / | 500/20 | 535/30 | 595/20 | 625/30 | 681/24 | 720/24 | / | 20x | / | 1 | 1 | 959 | 3 | no | 1 | 200 | 44 | 67.5 |
| S2 A | / | / | 480/20 | 520/40 | / | / | / | / | / | 12x water | / | 1 | 1 | 375 | 1 | no | 2 | 200 | 4 | 2.9 |
| S2 B | / | / | 480/20 | 520/40 | / | / | / | / | / | 12x | / | 1 | 1 | 811 | 4 | no | 1 | 200 | 18 | 25.4 |
| S2 C-D | 595/20 | 625/30 | 480/20 | 520/40 | / | / | / | / | / | 12x | / | 1 | 1 | 986 | 5 | no | 2 | 200 | 107 | 77.1 |
| S3 A | / | / | / | / | / | / | 681/24 | 720/24 | 2 | / | / | 2.5 | 2 | 393 | 2 | no | 1 | 200 | 8 | 12.2 |
| S3 B | / | / | / | / | / | / | 681/24 | 720/24 | 2 | / | / | 2.5 | 2 | 648 | 3 | no | 1 | 200 | 20 | 30.4 |
| M 1 | As in Fig. 2 A-A’’ | | | | | | | | | | | | | | | | | | | |
| M 2 | As in Fig. 2 B-B’’ | | | | | | | | | | | | | | | | | | | |
| M 3 | As in Fig. 3 A-C’’’ | | | | | | | | | | | | | | | | | | | |
| M 4 | / | / | / | / | 595/20 | 625/30 | 681/24 | 720/24 | / | 20x | / | 1 | 1 | 822 | 1 | no | 1 | 300 | 17 | 19.2 |
| M 5 | / | / | / | / | 595/20 | 625/30 | 681/24 | 720/24 | / | 20x | / | 2 | 1 | 256 | 1 | no | 1 | 300 | 3 | 4 |
| M 6 | As in Fig. 5 A-A’’ | | | | | | | | | | | | | | | | | | | |
