## Supplementary material for "A quick and versatile protocol for the 3D visualization of transgene expression across the whole body of larval *Drosophila*": Movie Legends

***Movie 1: Long-distance neuronal projections in the whole-body context: IR76b-Gal4 > UAS-mCherry-CAAX.***

Data set as used for Figure 2A-A’’, illustrating the traceability of sensory projections from the head region to the subesophageal zone as well as of v'td1 axonal projections from the abdominal segments to the central nervous system based on *IR76b-Gal4* driven mCherry-CAAX. Each time point represents a restricted volume (substack) in side view (rostral to the left). Grid spacing: 200 µm. The movie consists of 900 time points at a frame rate of 25 frames per second in HD 1280 x 720 (16:9) format.

***Movie 2: Long-distance neuronal projections in the whole-body context: nsyb-Gal4 > UAS-mCherry-CAAX.***

Data set as used for Figure 2B-B‘‘, showing the caudal region of a larva with pan-neuronally driven (*nsyb-Gal4*) mCherry-CAAX (white) overlaid with the autofluorescence signal (AF; green). Motor neuron projections (white) innervating the hindgut are presented as 3D views to allow better visualization. Caudal is to the bottom. Grid spacing: 100 µm. The movie consists of 300 time points at a frame rate of 25 frames per second in HD 1280 x 720 (16:9) format.

***Movie 3: Monomeric IFP as a superior clearing-resistant reporter: nsyb-Gal4 > UAS-mIFP-T2A-HO1.***

Data set as used for Figure 3A-C’’’, with pan-neuronally driven (*nsyb-Gal4*) mIFP. The movie starts from dorsal towards the central body region of the larva to visualize nerve fibers in distal regions, thereby highlighting neuromuscular junctions (NMJs) as an example. Turning towards rostral, the central nervous system (CNS) and nerve fiber projections become visible. During the movie, image contrast settings were adjusted according to the signal intensity of the respective structures. The movie consists of 600 time points at a frame rate of 25 frames per second in HD 1280 x 720 (16:9) format.

***Movie 4: Two-color fluorescence imaging in the whole-body context: Mb247:mCherry-CAAX + APL-Gal4 > UAS-mIFP-T2A-HO1.***

Close-up of a larval brain region, with mCherry-CAAX expressed in the mushroom bodies (by means of the MB247 promotor; magenta) and mIFP expressed in the APL neuron (by *APL-Gal4*, green). The movie shows the sites of innervation of the APL neurons within the calyx and lobes of the mushroom bodies. Grid spacing: 10 µm. Imaging parameters are given in Supplementary Table 1. The movie consists of 450 time points at a frame rate of 25 frames per second in HD 1280 x 720 (16:9) format.

***Movie 5: Two-color fluorescence imaging in the whole-body context: Mb247:mCherry-CAAX + IR76b-Gal4 > UAS-mIFP-T2A-HO1.***

Close-up of a larval brain region, with mCherry-CAAX expressed in the mushroom bodies (by means of the MB247 promotor; magenta) and mIFP expressed in sensory neurons (*by IR76b*-Gal4, green), showing the entrance of nerve fibers, originating from the head, into the subesophageal zone. Note that there is no overlap of the two populations of labeled neurons or their projections. Imaging parameters are given in Supplementary Table 1. Grid spacing: 20 µm. The movie consists of 300 time points at a frame rate of 25 frames per second in HD 1280 x 720 (16:9) format.

***Movie 6: Three-color fluorescence imaging in the whole-body context: PMCA$^{Venus\text{-}PT}$ + MB247:mCherry-CAAX + nsyb-Gal4 > UAS-mIFP-T2A-HO1.***

Data set as used for Figure 5A-A', with Venus-tagged PMCA encoded by the endogenous *PMCA* locus, mCherry-CAAX expressed in mushroom bodies by means of the MB247 promoter, and mIFP driven by *nsyb-Gal4*. The movie starts out showing the PMCA$^{Venus}$ signal (turquoise) from a dorsal view, then switches after 3 s to the mCherry-CAAX signal (magenta), and after another 3 s to the mIFP signal (yellow). Hereupon it zooms into the rostral region, showing the central nervous system and the afore-mentioned signals, this time in the opposite sequence. After 25 s the mCherry-CAAX signal in the mushroom bodies is shown (magenta), merged with the PMCA$^{Venus}$ signal (turquoise). Then the mCherry-CAAX signal in the mushroom bodies (magenta) is briefly shown merged with the mIFP-labeled nerve fibers (yellow) before the movie continues for only the mIFP signal towards the abdominal region, where neuromuscular junctions (NMJs) become visible. After a while, the PMCA$^{Venus}$ signal is added (turquoise) before a still framing shows the overlay of all three channels. During the movie, image contrast settings were adjusted according to the signal intensity of the respective structures. Grid spacing: 200 µm. The movie consists of 900 time points at a frame rate of 25 frames per second in HD 1280 x 720 (16:9) format.
